# *NEUROG2* regulates a human-specific neurodevelopmental gene regulatory program

**DOI:** 10.1101/2024.01.11.575174

**Authors:** Vorapin Chinchalongporn, Lakshmy Vasan, Fermisk Saleh, Dawn Zinyk, Hussein Ghazale, Ana-Maria Oproescu, Shruti Patel, Matthew Rozak, Yutaka Amemiya, Sisu Han, Alexandra Moffat, Sandra E Black, JoAnne McLaurin, Jamie Near, Arun Seth, Maged Goubran, Orly Reiner, Satoshi Okawa, Carol Schuurmans

**Affiliations:** Sunnybrook Research Institute, Biological Sciences Platform, Hurvitz Brain Sciences Program, 2075 Bayview Ave, Toronto, ON, Canada, M4N 3M5; Department of Biochemistry, 1 King’s College Cir, University of Toronto, ON, Canada, M5S 1A8; Department of Laboratory Medicine and Pathobiology, 1 King’s College Cir, University of Toronto, ON, Canada, M5S 1A8; Department of Medical Biophysics, 101 College St Suite 15-701, Toronto General Hospital, University of Toronto, ON, Canada, M5G 1L7; Sunnybrook Research Institute, Physical Sciences Platform, Hurvitz Brain Sciences Program, 2075 Bayview Ave, Toronto, ON, Canada, M4N 3M5; Dr. Sandra Black Centre for Brain Resilience & Recovery, LC Campbell Cognitive Neurology Unit, Sunnybrook Research Institute, Toronto, Ontario, Canada; Hurvitz Brain Sciences Program; Department of Medicine (Neurology) (SEB), University of Toronto, Toronto, Ontario, Canada; Departments of Molecular Genetics and Molecular Neuroscience, Weizmann Institute of Science, 76100, Rehovot, Israel; University of Pittsburgh School of Medicine, Vascular Medicine Institute, Department of Computational and Systems Biology, McGowan Institute for Regenerative Medicine, Pittsburgh, PA, USA

**Keywords:** cerebral organoid, CRISPR gene editing, lineage tracing, proneural gene, *NEUROG2*, extracellular matrix

## Abstract

Unique hallmarks of human neocortical development include slower rates of neurogenesis and the establishment of an extracellular matrix-rich, outer-subventricular zone that supports basal neural progenitor cell expansion. How gene regulatory networks have evolved to support these human-specific neurodevelopmental features is poorly understood. Mining single cell data from cerebral organoids and human fetal cortices, we found that *NEUROG2* expression is enriched in basal neural progenitor cells. To identify and purify *NEUROG2*-expressing cells and trace their short-term lineage, we engineered two *NEUROG2-mCherry* knock-in human embryonic stem cell lines to produce cerebral organoids. Transcriptomic profiling of mCherry-high organoid cells revealed elevated expression of *PPP1R17*, associated with a fast-evolving human-accelerated regulatory region, oligodendrocyte precursor cell and extracellular matrix-associated gene transcripts. Conversely, only neurogenic gene transcripts were enriched in mCherry-high cortical cells from *Neurog2:mCherry* knock-in mice. Finally, we show that *Neurog2* is sufficient to induce *Ppp1r17*, which slows human neural progenitor cell division, and *Col13a1*, an extracellular matrix gene, in P19 cells. *NEUROG2* thus regulates a human neurodevelopmental gene regulatory program implicated in supporting a pro-proliferative basal progenitor cell niche and tempering the neurogenic pace.

**SUMMARY STATEMENT:** Transcriptomic analyses of *NEUROG2-mCherry* knock-in human embryonic stem cell-derived cerebral organoids reveal a link between *NEUROG2* and extracellular matrix remodeling during human cortical development.

## INTRODUCTION

The rodent neocortex is smooth-surfaced, or lissencephalic, whereas, in larger mammals, including humans, neuronal layers are folded to create a gyrencephalic cortical structure marked by gyri and sulci. Understanding how these structural differences arise during development is an intensive area of investigation (Amin and Borrell, 2020; Moffat and Schuurmans, 2023). A key distinction between human and rodent corticogenesis is the evolution of unique neural progenitor cell (NPC) pools along with temporal differences in neuronal differentiation and maturation processes (Wallace and Pollen, 2023). Human NPCs have an overall longer neurogenic period, in part due to increased protein half-lives (Rayon et al., 2020) and longer cell cycle times, which are recapitulated in differentiation assays *in vitro* (Herculano-Houzel, 2012; van den Ameele et al., 2014). Additionally, a multitude of new genes have evolved to drive human brain development and function, originating either via gene duplications, structure/function alterations of ancestral genes or their gene regulatory regions, or from *de novo* origins (Hodge et al., 2019). Several human genes involved in neurodevelopmental processes are associated with fast-evolving regulatory regions, termed Human Accelerated Regions (HARs) (Girskis et al., 2021). One example of a critical HAR-regulated gene is *PPP1R17,* which is expressed in human cortical NPCs and not in rodent or ferret NPCs, and which lengthens the cell cycle when misexpressed in murine cortical cells (Girskis et al., 2021). However, it has yet to be determined how the expression of *PPP1R17* and other HAR-regulated genes are differentially regulated to support the unique developmental trajectory of human neocortices.

Initially, the cortical ventricular zone (VZ) is populated by primary NPCs, termed apical radial glia (aRG), which undergo direct neurogenesis to form deep-layer neurons. Later, aRG give rise to a secondary pool of intermediate progenitor cells (IPCs) that form a more basally-located subventricular zone (SVZ) (Vaid and Huttner, 2022). In rodent cortices, IPCs have a limited proliferative capacity and divide only 1-2 times before differentiating into late-born, upper-layer neurons (Haubensak et al., 2004; Miyata et al., 2004; Noctor et al., 2004). In contrast, in gyrencephalic species, NPCs have diversified and multiplied to drive cortical folding. In these folded cortices, early-born, deep-layer neurons are derived from aRG as well as from an expanded pool of basal radial glia (bRG) that lose their apical attachments and populate an enlarged outer subventricular zone (oSVZ) (Betizeau et al., 2013; Hansen et al., 2010; LaMonica et al., 2013; Martinez-Cerdeno et al., 2012; Shitamukai et al., 2011). Conversely, bRG are rare in rodent cortices, found mainly in dorsomedial domains (Vaid and Huttner, 2022) and representing only ∼5% of all NPCs (Martinez-Cerdeno et al., 2012; Shitamukai et al., 2011; Wang et al., 2011). In species with folded cortices, IPCs act as transit-amplifying cells, dividing multiple times before differentiating through a process termed indirect neurogenesis (Fietz et al., 2010). These rapidly expanding IPCs give rise to many more upper-layer neurons, with the increase in neuronal mass accommodated by the formation of cortical folds that expand the overall surface area (Fernandez and Borrell, 2023; Fietz et al., 2010). To support the expanded proliferative properties of basal progenitors in gyrencephalic species, basal NPCs produce an extracellular matrix (ECM)-rich niche that supports cell division (Amin and Borrell, 2020; Arai et al., 2011; Fietz et al., 2012; Florio et al., 2015; Martinez-Martinez et al., 2016; Pollen et al., 2015).

In invertebrate (e.g., *Drosophila*) and vertebrate (e.g., mouse and man) species alike, proneural genes encode basic-helix-loop-helix (bHLH) transcription factors that are evolutionarily-conserved drivers of neurogenesis (Oproescu et al., 2021; Quan et al., 2016). The proneural gene *Neurog2* is the main driver of murine neocortical development. In the developing rodent neocortex, *Neurog2* is required and sufficient to specify a glutamatergic neuronal identity (Fode et al., 2000; Mattar et al., 2008; Oproescu et al., 2021; Schuurmans et al., 2004). The related gene, *Neurog1*, also specifies a glutamatergic neuronal fate in the developing neocortex, but functions at a slower rate, and tempers the pace of early cortical neurogenesis by heterodimerizing with *Neurog2* to suppress its potent neurogenic activity (Han et al., 2018).

Due to the inaccessibility of fetal brain tissue, uncovering human-specific features of proneural gene function during human neocortical development is a major challenge. To circumvent these issues, 3D brain organoids can be generated from human embryonic stem cells (hESCs) or induced pluripotent stem cells (iPSCs). These organoids can be directed to differentiate into specific neural tissues, such as the cerebral cortex, optic vesicles, midbrain, spinal cord and choroid plexus using activators and inhibitors of signaling pathways that drive normal central nervous system development (Amin et al., 2023; Karzbrun and Reiner, 2019). Cerebral organoids are advantageous as they can be grown for a year or more *in vitro* (Lancaster et al., 2013; Qian et al., 2016) or transplanted in rodent brains for further maturation (Revah et al., 2022; Wilson et al., 2022). Further features of cerebral organoids that show a resemblance to the neocortex; they are comprised of neural progenitor cells (NPCs) that differentiate into glutamatergic and GABAergic neurons, astrocytes and oligodendrocytes (Karzbrun and Reiner, 2019; Lancaster and Knoblich, 2014; Lancaster et al., 2013; Qian et al., 2016)

Here, we have begun to dissect the role of *NEUROG2* in human cortical neurogenesis using a cerebral organoid model system. We found that *NEUROG2* is expressed in human fetal cortices and in cerebral organoids, with the highest transcript counts detected in IPCs and aRG, followed by smaller subsets of bRG and neurons. By engineering *NEUROG2-mCherry* knock-in (KI) hESC lines, and creating cerebral organoids from these cells, we were able to compare the transcriptomes of mCherry-high versus mCherry-low cells. Unexpectedly, mCherry-high cells were enriched in oligodendrocyte precursor cell (OPC) and ECM-related transcripts, contrasting to the *Neurog2-mCherry*-high lineage in rodent cortices, in which neurogenic gene expression was instead elevated. *PPP1R17* transcripts were also enriched in mCherry-high cells in *NEUROG2-mCherry* KI-hESC-derived organoids, which is of interest given the link between *PPP1R17* and human neocortical evolution (Girskis et al., 2021). Finally, *Neurog2* was sufficient to induce the expression of *Ppp1r17* and *Col3a1*, an ECM gene, in P19 cells. Given the significance of the ECM to basal progenitor cell expansion (Amin and Borrell, 2020; Arai et al., 2011; Fietz et al., 2012; Florio et al., 2015; Martinez-Martinez et al., 2016; Pollen et al., 2015), and human *PPP1R17* to slowing down cell cycle progression and overall rates of neurogenesis (Girskis et al., 2021), our study highlights important ways in which *NEUROG2* function has adapted to regulate human-specific features of neocortical development.

## RESULTS

### *NEUROG2* is the predominant proneural gene in human fetal cortices and cerebral organoids

In rodent cortices, *Neurog2* is the predominant proneural gene, whereas the related gene *Neurog1* is expressed at lower levels and in a shorter time window (Han et al., 2018; Moffat et al., 2023). To determine whether *NEUROG2* similarly predominates during human cortical development, we assayed *NEUROG1* and *NEUROG2* expression profiles in existing scRNA-seq data from human fetal cortical cells collected between gestational week (GW) 08 to GW26, during the period of active neurogenesis (Zhong et al., 2018). Of the cells assigned an NPC identity, the majority expressed *NEUROG2* (72.06%), either together with *NEUROG1* (38.62%) or alone (33.4%) (Fig. 1A). In contrast, *NEUROG1* was expressed in fewer human fetal cortical cells (47.93%) and only 9.31% of cortical NPCs expressed *NEUROG1* alone (Fig. 1A). The expression of *NEUROG1* and *NEUROG2* remained elevated from GW 09 and persisted until GW 26 (Fig. 1A), contrasting to the reduction in *Neurog1* expression and double^+^ cells observed at the midpoint (*i.e.,* embryonic day 15.5) of murine cortical development (Han et al., 2018).

**Figure 1.**
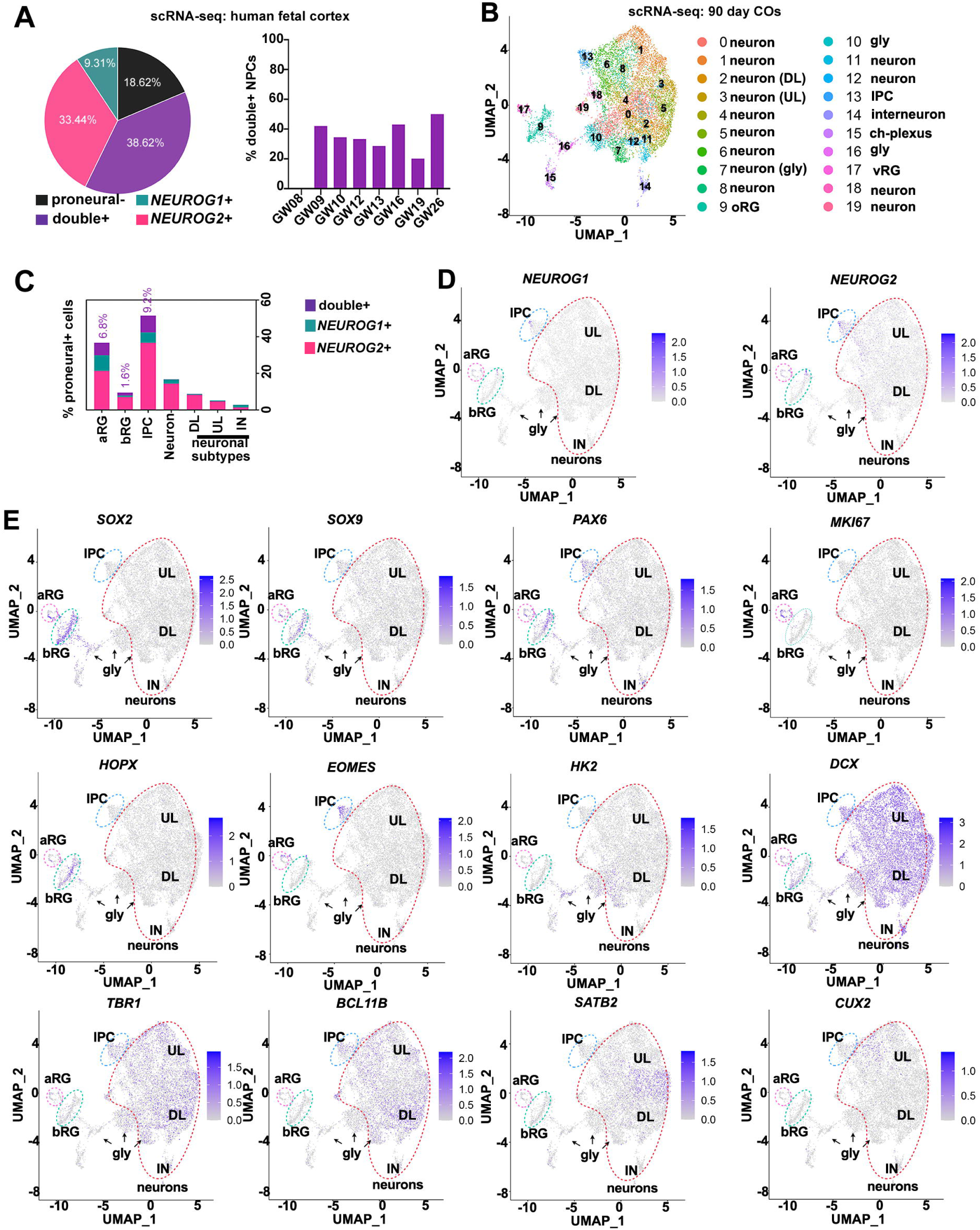
Transcriptomic assessment of *NEUROG1* and *NEUROG2* expression in gestational week 8-26 human fetal cortices and 3-mo old cerebral organoids. (A) Distribution of *NEUROG1/NEUROG2* single and double^+^ cells in scRNA-seq datasets from human fetal cortices between gestational week 8 to 26. (B-E) UMAP of scRNA-seq data from 3-month-old cerebral organoids (B), showing proportions of cortical cell types expressing *NEUROG1* and/or *NEUROG2* (C). Key genes made for cluster assignments are mapped onto the UMAP, including *NEUROG1* and *NEUROG2* (D) and cell type-specific markers (E). aRG, apical radial glia; bRG, basal radial glia; ch-plexus, choroid plexus; DL, deep layer; Gly, high glycolytic cells; IN, interneuron; IPC, intermediate progenitor cell; UL, upper layer.

We circumvented the inaccessibility of fetal brain tissues by using human cerebral organoids, as a surrogate model of human corticogenesis, to assay proneural gene function. To confirm that hESC-derived cerebral organoids expressed *NEUROG1* and *NEUROG2*, and to identify expressing cell types, we mined scRNA-seq data from 90-day old cerebral organoids generated with a Lancaster protocol (Sivitilli et al., 2020). Unbiased Seurat clustering produced 19 cell clusters, which were then stratified into distinct cell types based on the expression of cell-type specific markers (Fig. 1B-E, Table S1), as previously described (Sivitilli et al., 2020). We found that *NEUROG1* and *NEUROG2* expression was highest in IPCs, of which, more than 50% expressed one of these proneural genes, with *NEUROG2* transcripts detected in more cells than *NEUROG1* mRNA (Fig. 1C-E). *NEUROG1* and/or *NEUROG2* transcripts were also detected in ∼40% of aRG and ∼10% of bRG, with *NEUROG2* transcripts similarly more frequently detected in these cells compared to *NEUROG1* mRNA (Fig. 1C-E). Finally, we also detected *NEUROG2* transcripts in a small percentage of cortical neurons, whereas *NEUROG1* expression was nearly absent in neuronal populations (Fig. 1C-E). Thus, *NEUROG2* expression is more widespread among human fetal cortical cells and human cerebral organoid cells than *NEUROG1*.

To understand the lineage dynamics associated with human proneural gene expression during corticogenesis, we computationally isolated cortical cells expressing *NEUROG1* and/or *NEUROG2* from scRNA-seq data of 90-day-old human cerebral organoids (Sivitilli et al., 2020) and generated pseudotime trajectories. The resultant pseudotime ordering of cells along a lineage trajectory revealed a single branch point and three cell states (Fig. 2A,B). State 1 cells had the earliest pseudotime identity and included the highest fraction of *NEUROG1/NEUROG2* double*^+^* and *NEUROG1* single^+^ cells, and the lowest fraction of *NEUROG2* single^+^ cells (Fig. 2B-D). We determined that state 1 cells had the highest levels of aRG, bRG, and proliferating cell-associated transcripts by mapping the expression of cell identity markers onto the lineage trajectory (Fig. 2E, F; Fig. S1A; Table S1). In state 2 cells, which had an intermediate pseudotime identity, *NEUROG1* expression and *NEUROG1/NEUROG2* co-expression declined, while the fraction of *NEUROG2* single^+^ cells increased slightly (Fig. 2B-D). In state 2 cells, aRG/bRG-marker expression declined, while there was an increase in IPC markers, especially in *NEUROG2* single^+^ cells (Fig. 2E, F). Finally, state 3 cells, which had the latest pseudotime identities, predominantly expressed *NEUROG2* alone, with very low fractions of the *NEUROG1/NEUROG2* double^+^ and *NEUROG1* single^+^ pools (Fig. 2B-D). State 3 cells displayed elevated levels of early neuronal marker transcripts, including deep-layer 5 and 6 markers (Fig. 2B-F, Table S1). Thus, *NEUROG1* expression is confined to early stages of neural lineage development in human cerebral organoids, and is enriched in aRG and bRG. In contrast, while *NEUROG2* is also expressed in aRG and bRG, the proportion of *NEUROG2-*expressing cortical cells increases as they mature into IPCs, and transcripts are also detected in neurons in later pseudotime states 2 and 3.

**Figure 2.**
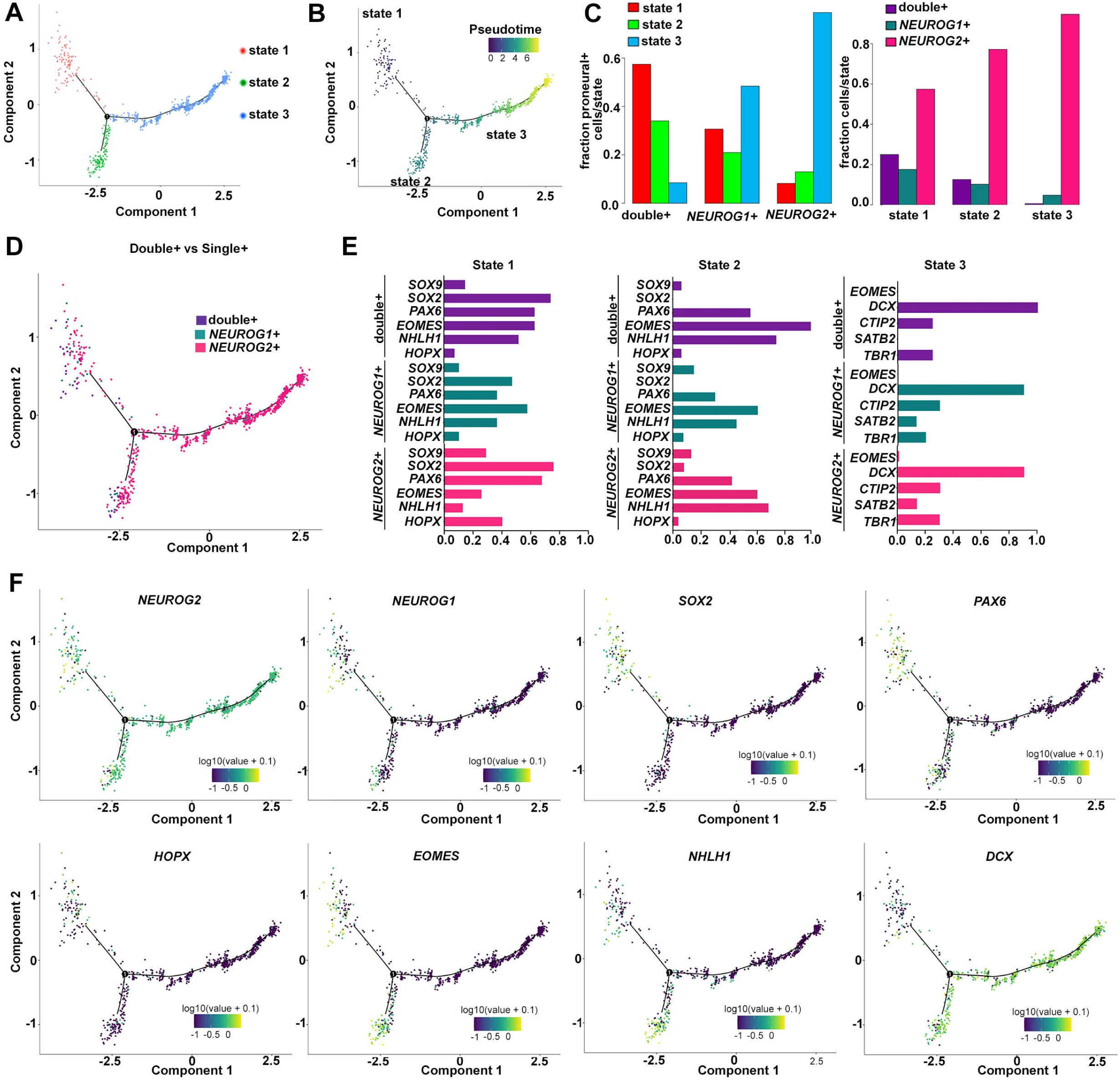
Pseudotime trajectory analysis of *NEUROG1* and *NEUROG2* expressing cerebral organoid cells. (A-F) Monocle3 lineage trajectory analysis of *NEUROG2*^+^, *NEUROG1*^+^ and double^+^ cells in 3-mo cerebral organoid scRNA-seq data (Sivitilli et al., 2020) (A), showing a pseudotime trajectory (B), and the distributions of proneural-expressing cells in states 1, 2 and 3 *NEUROG1* and/or *NEUROG2* expressing cells were mapped onto the pseudotime trajectory (D), along with other key markers of cortical cell types (F). The distributions of proneural-expressing cells in each cell state expressing cell type-specific markers were plotted (E).

### CRISPR engineering to generate *NEUROG2-mCherry-*KI hESCs for cerebral organoid production

For the remainder of this study, we focused on *NEUROG2*, given its predominant expression in human cerebral lineages. To isolate and characterize *NEUROG2-*expressing cells from cerebral organoids, we used CRISPR/Cas9 to engineer *NEUROG2-mCherry* KI reporter hESC lines (Fig. S2A,B). Individual clones were screened for homology-directed repair using digital droplet PCR (Fig. S2B,C). Of the 87 clones that were selected for screening, 4 clones had a correctly targeted *NEUROG2-mCherry* insertion (4.6% target efficiency). To ensure that the gene targeting process did not introduce any unintended mutations in the *NEUROG2* locus, we sequenced the knock-in clones at the 5’ and 3’ junction of the Cas9 targeted site. No mutations were detected in the two correctly targeted clones, which we designated as lines #105 and #117 (Fig. S2D). In addition, both *NEUROG2*-mCherry KI hESC lines expressed pluripotency genes (*OCT4, SOX2, NANOG*), and did not show genomic abnormalities in the most common mutated regions reported in hESCs (Fig. S3A-C). We thus followed through with the two correctly targeted *NEUROG2-mCherry* KI hESC clones (#105, #117) for downstream experiments.

To investigate *NEUROG2* function, we employed a modified Lancaster protocol incorporating dual Smad inhibitors to generate hESC-derived cerebral organoids that are enriched in cortical gene expression (Qian et al., 2018) (Fig. 3A,B). To confirm neural induction and organoid maturation with this approach, cerebral organoids were isolated on day 42, optically cleared, co-immunostained with SOX2 to label NPCs and TUJ1 (b3-tubulin) to mark newborn neurons, and imaged in wholemount using lightsheet microscopy (Fig. 3C). 3D imaging revealed a network of TUJ1^+^ axonal fibers encasing the 42-day old cerebral organoids. In some instances, neural rosettes comprised of an inner SOX2^+^ NPC layer and outer TUJ1^+^ neuronal layer appeared as external protrusions (Fig. 3C). In 2D sections, day 30-35 organoids contained neural rosettes that expressed the pan-NPC markers SOX2 and NESTIN, as well as PAX6, a regional marker of a cortical NPC identity (Fig. 3D). Newborn neuronal markers, such as DCX and TUJ1, labeled neurons and their processes on the outside of the neural rosettes (Fig. 3D).

**Figure 3.**
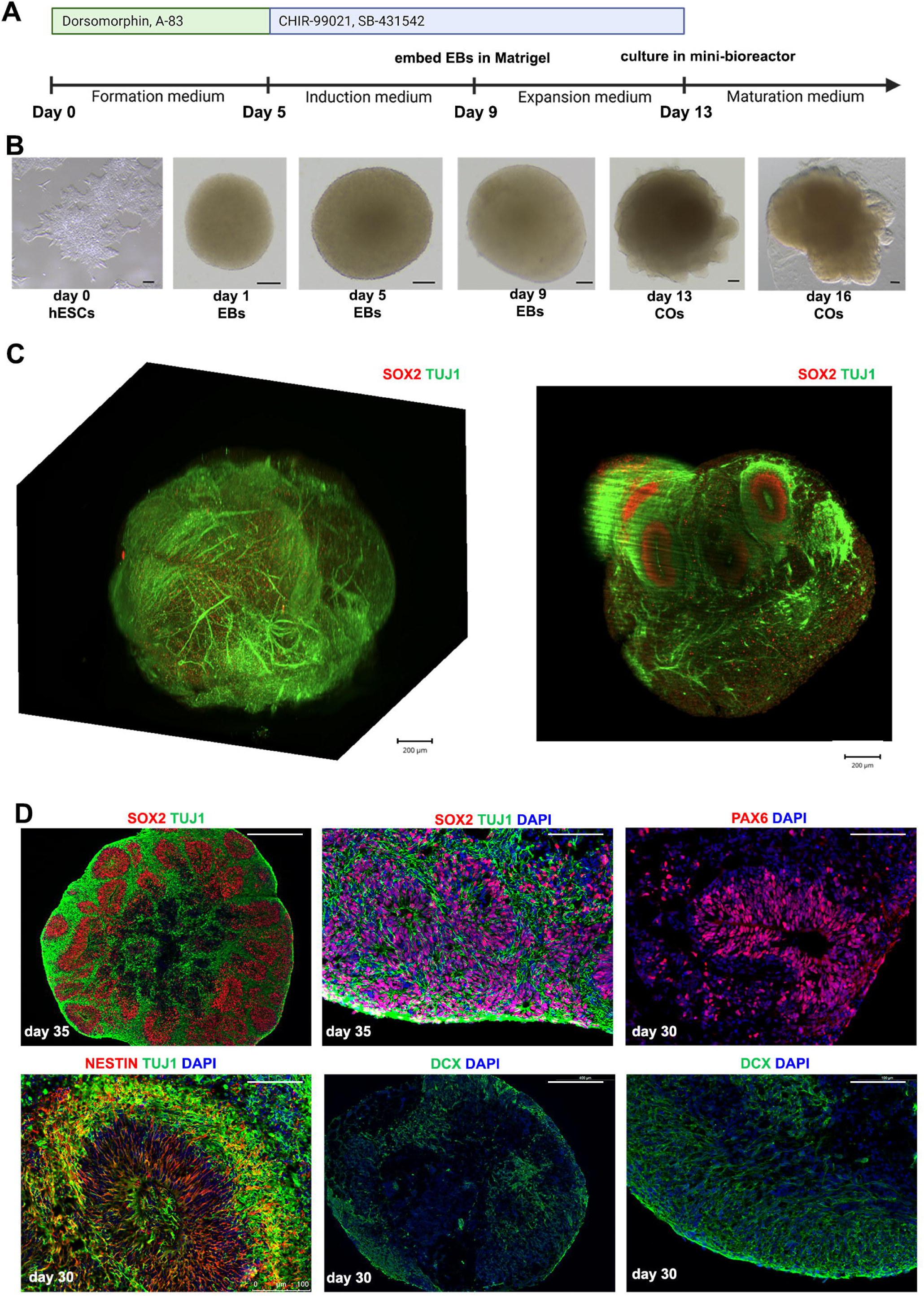
Generation of hESC-derived cerebral organoids using dual Smad inhibition. (A,B) Experimental protocol used for cerebral organoid generation (A) and photomicrographs of derivative cells and cerebral organoids at various stages of cerebral organoids formation (B). (C) 3D light-sheet imaging of two-day 42 cerebral organoids that were cleared and immunolabeled for SOX2 (red) and TUJ1 (green). (D) Immunostaining of cryosections of day 30-35 cerebral organoids immunostained for SOX2/TUJ1, PAX6, NESTIN/TUJ1 and DCX. Scale bars in B, 100 µm; in C, 200 µm; in D, 400 µm in low magnification images and 100 µm in high magnification images.

To confirm the competence of *NEUROG2-mCherry* KI hESC clones to generate cerebral organoids, we followed the same dual SMAD inhibitor protocol. Notably, mCherry expression was more widespread than NEUROG2 due to the inherent characteristics of the proteins involved: mCherry is a stable protein with a long intracellular half-life, while NEUROG2 has a relatively short half-life of approximately 30 minutes (Li et al., 2012). The long half-life of mCherry makes it suitable for lineage tracing of cells derived from NEUROG2^+^ NPCs, at least in the short-term. In day 18 organoids, we detected robust co-expression of NEUROG2 and mCherry in neural rosettes encircling the periphery of the organoid (Fig. 4A). mCherry was co-expressed with additional NPC markers, including SOX2 (Fig. 4B) and PAX6 (Fig. 4C), consistent with a neural and cortical NPC identity, respectively. By day 45, SOX2 expression remained widespread in neural rosettes, whereas NEUROG2 and PAX6 were expressed in relatively fewer cells by this stage of organoid development (Fig. 4D-F). In contrast, mCherry expression persisted in scattered rosettes throughout the day 45 organoids, albeit with limited co-localization between mCherry and NPC markers (Fig. 4D-F). In the BrainSpan Atlas, *NEUROG2* transcript levels similarly decline between GW 9 and 12 in human fetal cortices and are reduced over time in cortical brain organoids (Cheroni et al., 2022). Overall, these data demonstrate that mCherry expression persists after *NEUROG2* expression declines, allowing us to use mCherry to profile the phenotype of organoid cells derived from *NEUROG2*^+^ NPCs.

**Figure 4.**
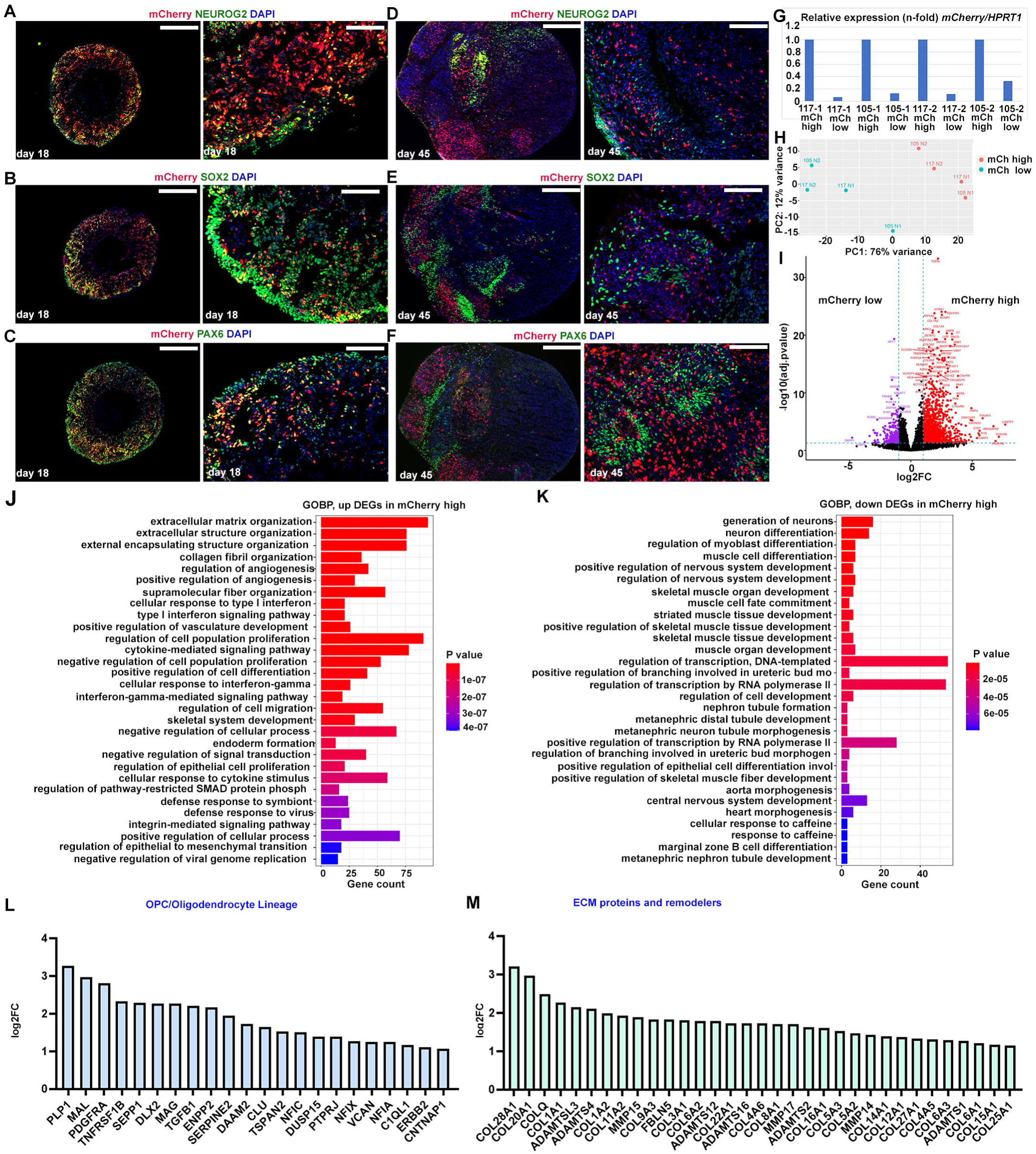
Analyses of mCherry-high and mCherry-low cells in *NEUROG2-mCherry* KI hESC-derived cerebral organoids. (A-F) Immunostaining of NEUROG2-mCherry KI hESC-derived cerebral organoids collected at day 18 (A-C) and day 45 (D-F) and analysed for the expression of mCherry with NEUROG2, SOX2 and PAX6. (G) qPCR to validate FACS-enrichment of mCherry (mCh)-high and mCh-low cells collected from two *NEUROG2* mCherry KI hESC cell lines (#105 and #117) from different sets of cerebral organoids (−1, −2). (H-K) PCA analysis of bulk RNA-seq data collected from mCherry-high and mCherry-low cells (H). Volcano plot showing an enrichment of more genes in mCherry-high versus mCherry-low cells Differentially expressed genes (DEGs) in mCherry-high cells were identified compared to mCherry-low cells (I). Biological Process Gene Ontology (GO) terms enriched in DEGs that were upregulated (J) or downregulated (K) in mCherry-high cells. (L-M) Bar graphs showing the log2FC values of the differentially expressed genes, OPC lineage markers (L) and ECM proteins and remodelers (M). DEGs, differentially expressed genes; ECM, extracellular matrix; mCh, mCherry; OPC, oligodendrocyte precursor cells.

### Transcriptomic analyses reveal a link between *NEUROG2*, the ECM and oligodendrocyte precursor cells

To examine the molecular phenotype of cells in the *NEUROG2*^+^ lineage, cerebral organoids were generated from two *NEUROG2-mCherry* KI hESC lines (#105 and #117) in two independent experiments. Cerebral organoids were harvested after 45-47 days *in vitro*, at a stage when cerebral organoids, generated with the Qian et al. (2016, 2018) protocol, share a transcriptional signature with GW 9-12 fetal cortices, according to the BrainSpan Atlas (Cheroni et al., 2022). mCherry-high and mCherry-low cells were isolated by FACS from pools of 7-8 cerebral organoids and the enrichment of *mCherry* transcripts in mCherry-high versus -low cells was verified by qPCR (Fig. 4G). To profile gene expression, we used targeted transcriptome sequencing with a human gene expression assay that covers 20,802 human genes, representing >95% of the UCSC reference genome (Table S2). Principal component analysis (PCA) of gene expression datasets revealed that mCherry-high versus mCherry-low cells were transcriptionally divergent and segregated from each other irrespective of the initiating cell line or experimental day (Fig. 4H). A comparative analysis of differentially expressed genes (DEGs) in mCherry-high versus mCherry-low cells identified 1204 genes enriched in mCherry-high cells and 263 genes enriched in mCherry-low cells (Fig. 4I, Table S3). Biological Process gene ontology (GO) analysis of DEGs revealed an enrichment of terms associated with the extracellular matrix (ECM) in mCherry-high cells, including ‘*extracellular matrix organization’* and ‘*collagen fibril organization’* (Fig. 4J; Table S4). Conversely, neuronal differentiation GO terms such as ‘*generation of neurons’* and ‘*neuron differentiation’* were over-represented in mCherry-low cells (Fig. 4K; Table S5).

To characterize the two populations of cerebral organoid cells more thoroughly, we selectively assayed transcript counts for known genes involved in cortical development, focusing on transcripts with a log-2-fold-change (log2FC) greater than 1 (*i.e.,* a doubling in the original scaling) and with a padj<0.05, corresponding to a false discovery rate cutoff of 0.05. Markers of apical RG, basal RG and proliferating cells were expressed at relatively equivalent levels in mCherry-high and mCherry-low cerebral organoid cells (Fig. S4A-C, Table S2). Of the previously characterized human IPC markers, only *PPP1R17* (Pebworth et al., 2021) and *NHLH1* were expressed at detectable levels in day 45-47 cerebral organoids, and strikingly, there was a 3.2 log2FC in *PPP1R17* transcripts, which were elevated in mCherry-high cells (Fig. S4D). Thus, *NEUROG2-mCherry* KI hESC-derived cerebral organoids are comprised of both mCherry-high and low cells that express relatively equivalent levels of aRG, bRG and IPC markers, with the exception of an enrichment in *PPP1R17* gene transcripts in mCherry-high IPCs.

We next assessed transcript levels for pan-neuronal markers in the two cerebral organoid populations to characterize lineage preference (Table S2). Of these pan-neuronal markers, only *GAP43* was enriched in mCherry-high versus mCherry-low cerebral organoid cells at day 45-47 (Fig. S4E). In addition, transcripts for two glutamatergic-specific neuronal genes, *SLC17A6/VGLUT2* (1.03 log2FC) and *SLC17A7/VGLUT1* (2.36 log2FC), were more abundant in mCherry-high cerebral organoid cells (Fig. S4F). This finding was anticipated given that *Neurog2* is necessary and sufficient to specify a glutamatergic neuronal fate (Dixit et al., 2014; Li et al., 2012; Schuurmans et al., 2004). Unexpectedly, *NEUROD4* and *NEUROD6*, which act downstream of *NEUROG2* in a neurogenic cascade, were slightly enriched in mCherry-low cells, possibly due to the reduction in mCherry expression as NPCs differentiate (Fig. S4F). In addition, two GABAergic neural lineage markers (e.g., *ASCL1, GAD2*) were elevated in the mCherry-low lineage (Fig. S4G). These findings are in keeping with the known cross-repressive interactions between *Neurog2* and *Ascl1,* a different proneural gene that specifies GABAergic neuronal fate in cortical NPCs (Han et al., 2021; Schuurmans et al., 2004).

Next, we examined layer 1-6 neuronal markers to conclude our characterization of neuronal diversity in the cerebral organoid populations (Fig. S4I, Table S2). There were relatively equal transcript levels for most laminar markers in mCherry-high and mCherry-low cerebral organoid cells, except for increased transcript counts for the layer 5 marker *PCP4* (1.68 log2FC) and the layer 1 marker *RELN* (2.05 log2FC) in mCherry-high cerebral organoid cells. Conversely, the layer 5 marker *BCL11B* (1.45 log2FC) and layer 6 marker *FOXP2* (1.21 log2FC) transcript counts were higher in mCherry-low cells (Fig. S4I). These data suggest that there may be unique laminar biases for neurons derived from *NEUROG2* positive versus negative NPCs.

Lastly, we examined the representation of glial cells amongst the mCherry-high and mCherry-low populations. We detected transcripts for several astroglial cell markers at equivalent levels in mCherry-high versus low cells (Fig. S4H). In contrast, 28/221 genes expressed in OPCs and/or involved in oligodendrocyte development (GO:0048709) were among the DEGs that were upregulated in mCherry-high cerebral organoid cells (Fig. 4L, Table S6). Taken together, these data reveal an unexpected enrichment of *PPP1R17*, an IPC-expressed gene that is associated with a fast-evolving human-accelerated regulatory region, and OPC-related genes in *NEUROG2-mCherry* KI-high cerebral organoid cells.

### ECM protein deposition adjacent to mCherry^+^ cells in *NEUROG2-mCherry* KI hESC-derived cerebral organoids

A pro-proliferative ECM niche that supports integrin signaling is produced by basal NPCs in human and not in rodent cortices (Amin and Borrell, 2020; Arai et al., 2011; Fietz et al., 2012; Florio et al., 2015; Martinez-Martinez et al., 2016; Pollen et al., 2015). However, even though ECM genes are transcribed in germinal compartments in human fetal cortices (Fietz et al., 2012), ECM proteins are deposited in the cortical plate, such that transcripts and proteins are not detected in the same compartments (Long et al., 2018). Given the enrichment of GO terms associated with the ECM in *NEUROG2-mCherry* KI-high cerebral organoid cells, we further mined transcript read count data for ECM-related genes from GO: 0030198, of which 76/327 genes were among the DEGs (Table S7). ECM genes that were expressed at elevated levels by mCherry-high cerebral organoid cells included collagens, and ADAM and matrix metalloproteases, which are involved in ECM remodeling (Fig. 4M, Table S2).

To examine whether ECM proteins were detected in *NEUROG2-mCherry* KI hESC-derived cerebral organoids, we performed immunostaining with COL4 and LAM. In day 18 cerebral organoids, robust expression of COL4 and LAM was detected in circular formations surrounding the necrotic core, as well as in protrusions into the organoid periphery, where mCherry^+^ cells were located (Fig. 5A). In higher magnification images, there was limited overlap between COL4 or LAM and mCherry (Fig. 5A), which was not necessarily surprising given that ECM proteins are produced by NPCs but finally deposited in the cortical plate. At day 45, there was a change in the distribution of ECM proteins in *NEUROG2-mCherry* KI hESC-derived cerebral organoids. At this stage, COL4^+^ and LAM^+^ fibrils invaded the neural rosettes, and LAM was enriched in the periphery of the organoids (Fig. 5B). A necrotic core was no longer visible at this stage (Fig. 5B). To better visualize the distribution of ECM proteins at a later stage, tissue clearing, immunolabeling, and lightsheet fluorescence microscopy was performed on day 119 organoids. In these cleared organoids, mCherry^+^ cells were scattered throughout the organoid interior, COL4 was similarly detected in the center of the organoids, and LAM proteins encased the outer surface of the organoid (Fig. 5C,D).

**Figure 5.**
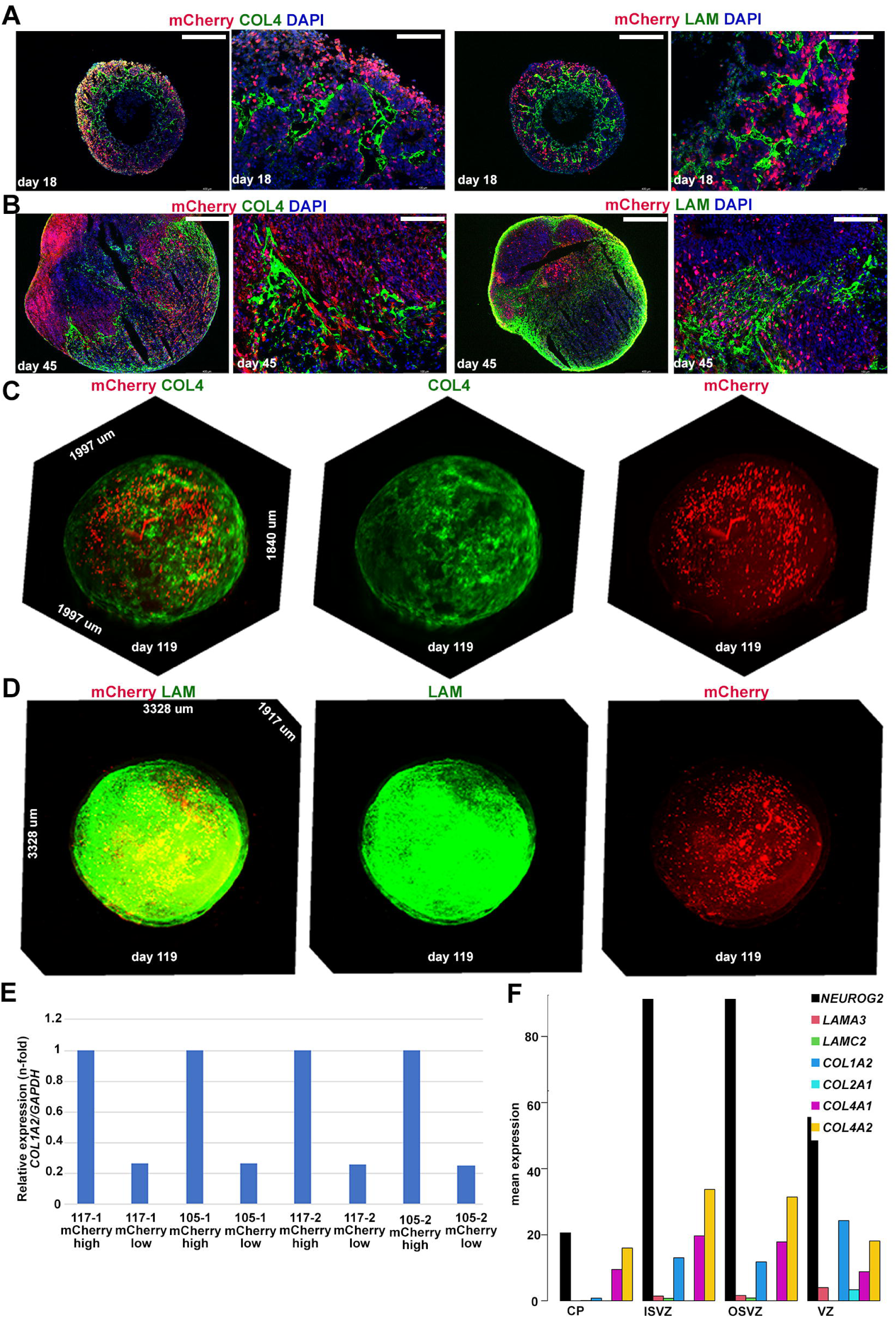
Expression of ECM markers in *NEUROG2*-mCherry KI hESC derived cerebral organoid. (A,B) Expression of mCherry with ECM markers Collagen IV (COL 4) and Laminin (LAM) in day 18 (A) and day 45 (B) *NEUROG2*-*mCherry* derived cerebral organoids. (C,D) 3D light-sheet imaging of two-clarified day 119 cerebral organoids immunolabeled for mCherry (red) and COL4 (green, C) or LAM (green, D). (E) ddPCR to look at the relative expression of COL1A2/GAPDH in mCherry-high versus mCherry-low cells isolated from day 45 cerebral organoids derived from the two *NEUROG2-mCherry* KI cell lines (#117 and #105). (F) Mining a bulk RNA-seq dataset from human fetal cortical zones (Fietz *et al*., 2012), revealed an enrichment of *NEUROG2* and ECM gene transcripts in the same cell populations.

Since the enrichment of ECM gene expression in mCherry-high cerebral organoid cells was unexpected, we confirmed this enrichment by performing qPCR on the FACS-enriched mCherry-high and -low cells, focusing on *COL1A2* (Fig. 5E). To independently validate the significance of the association between *NEUROG2* and ECM gene expression, we mined a bulk RNA-seq dataset from a study in which the human fetal cortex was dissected into germinal zones, leading to the conclusion that ECM genes support the proliferation of human NPCs (Fietz et al., 2012). In this dataset, several collagen genes, especially *COL1A2, COL4A1,* and *COL4A2*, were expressed at elevated levels in the VZ as well as inner and outer SVZ, in compartments in which *NEUROG2* transcript levels were also elevated (Fig. 5F). In conclusion, while mCherry^+^ cells do not directly co-localize with the ECM proteins, *NEUROG2*-expressing NPCs transcribe ECM genes that are then translated into proteins that are deposited throughout the cerebral organoid.

### *Neurog2* does not lineage trace ECM or OPC genes in the early embryonic murine cortex

The unforeseen enrichment of OPC and ECM genes in *NEUROG2*-mCherry-high organoid cells prompted us to compare to murine *Neurog2*-mCherry cortical lineages. We previously generated a *Neurog2-mCherry* KI transgenic line, from which we performed RNA-seq on mCherry-high versus proneural gene-‘negative’ NPCs collected from embryonic day (E) 12.5 cortices (Fig. 6A) (Han et al., 2021). E12.5 murine cortices are roughly at the same stage as GW10 human cortices, and have the equivalent of a day 45-47 cerebral organoid gene signature (Cheroni et al., 2022). ‘Proneural-negative’ NPCs also did not express Ascl1, which were negatively selected against using GFP expression from E12.5 *Neurog2-mCherry;Ascl1-GFP* double KI transgenic mice (Han et al., 2021). FACS was used to select mCherry^+^ cells that co-expressed CD15, an NPC marker, and ‘proneural-negative’ NPCs that expressed CD15 without mCherry or GFP expression were also collected (Han et al., 2021).

**Figure 6.**
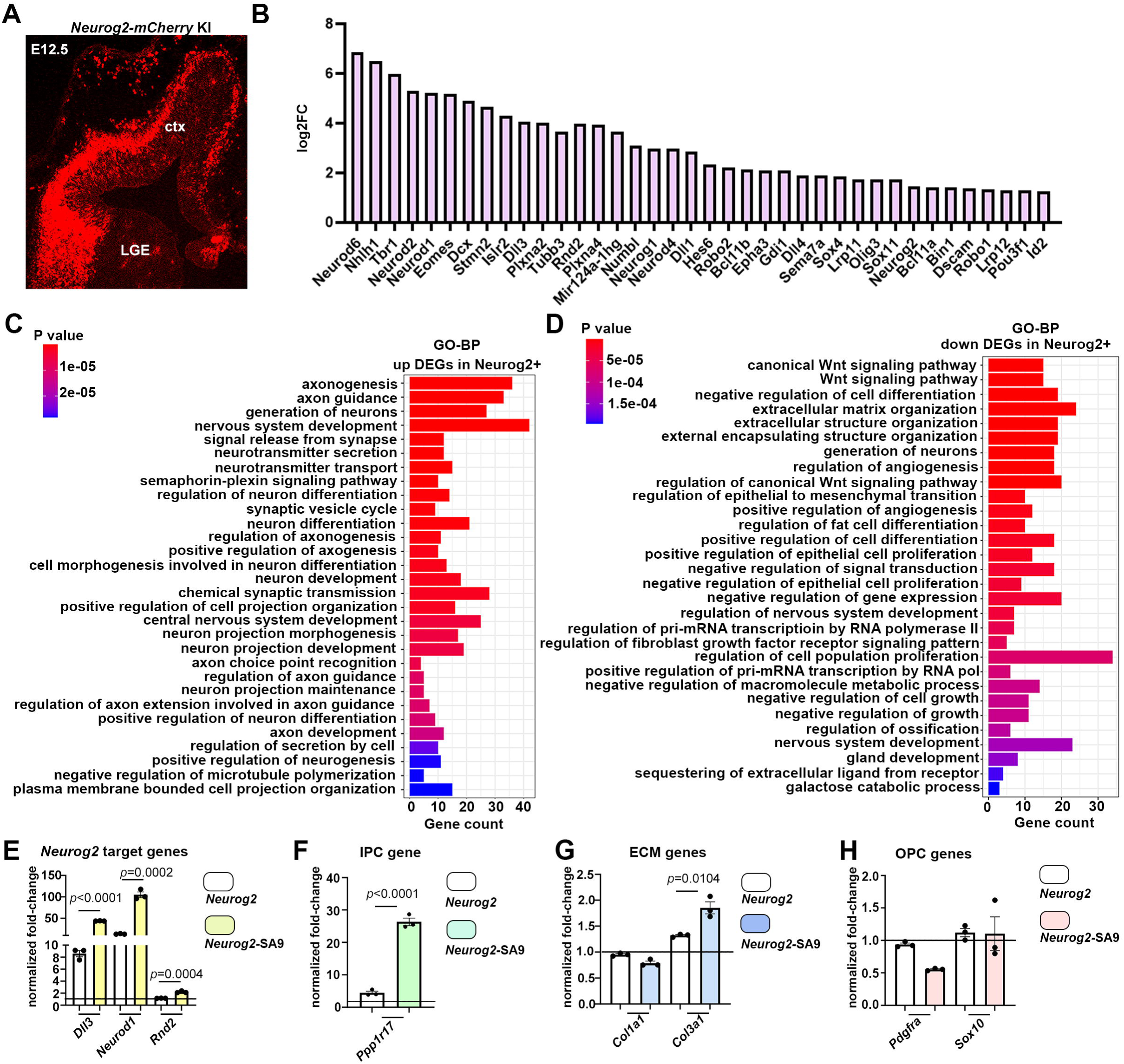
Characterization of mCherry positive cells in embryonic *Neurog2*-mCherry KI cortices. (A) mCherry expression in E12.5 *Neurog2-mCherry* KI mouse brain. (B) Transcriptomic analysis to compare the log2FC values of cell type specific markers in mCherry-high, CD15^+^ cortical NPCs to mCherry low cells (data from (Han et al., 2021)). Biological Process GO terms enriched in DEGs that were upregulated (C) or downregulated (D) in mCherry-high cells. (E-H) P19 cells transfected with pCIG2, pCIG2-*Neurog2* or pCIG2-*Neurog2-SA9* and analysed for the expression of known target genes (E), OPC genes (F), ECM genes (G) and IPC marker (H). Student t-tests were used to individually compare expression levels. p-values: ns -not significant, <0.05 *, <0.01 **, <0.001 ***. Significance was defined as p-values less than 0.05. DEGs, differentially-expressed genes; ECM, extracellular matrix; IPC, intermediate progenitor cell; mCh, mCherry; OPC, oligodendrocyte precursor cells.

Comparisons of bulk RNA-seq datasets from mCherry-high and proneural negative NPCs identified 1112 DEGs (padj<0.05; Table S8). Within the “*positive regulation of neurogenesis*” GO: 0050769 category, 61/329 genes were DEGs when comparing mCherry-high to proneural negative cortical cells (Fig. 6B; Table S9). Further GO term enrichment analysis identified multiple “Biological Process” GO terms associated with neurogenesis that were enriched in *Neurog2-mCherry* high cortical NPCs, including “*axonogenesis”,* “*generation of neurons”*, and “*nervous system development”* (Fig. 6C). Conversely, “*canonical Wnt signaling”* and “*extracellular matrix organization”* GO terms were under-represented in *Neurog2-mCherry*-high cortical NPCs (Fig. 6D), which is the opposite of the *NEUROG2-mCherry* data collected from cerebral organoids. Moreover, neither Collagen nor Laminin expression was detected in E14.5 *Neurog2-mCherry* KI cortices, despite their enrichment in the basal lamina at the pial surface (Fig. S6A-C). These data suggest that the association between *NEUROG2* and ECM gene expression is unique to human cortical lineages and are consistent with the finding that ECM production by human basal NPCs is a newly evolved driver of their increased proliferative potential (Amin and Borrell, 2020; Arai et al., 2011; Fietz et al., 2012; Florio et al., 2015; Martinez-Martinez et al., 2016; Pollen et al., 2015).

To further characterize *Neurog2*-expressing lineages in the early embryonic murine cortex, we next examined whether known determinants of murine neural cell fates and drivers of cortical neurogenesis were amongst the DEGs. A subset of aRG markers, including *Fabp7* (1.6 log2FC), *Hes1* (2.03 log2FC) and *Gli3* (1.26 log2FC), were enriched in *Neurog2-mCherry*-negative cortical NPCs (Fig. S5A). These data are consistent with the idea that *Neurog2* expression is turning on just as aRG transition to IPCs (Moffat et al., 2023), and/or suggest that there is a *Neurog2-*negative aRG lineage. Conversely, mCherry-high NPCs were enriched in the expression of IPC markers, including *Nhlh1* (6.48 log2FC), *Neurod1* (5.21 log2FC) and *Eomes* (5.19 log2FC) (Fig. S5B). Notably, *Ppp1r17* transcripts, which were elevated in mCherry-high cerebral organoid cells, and mark human IPCs, were not detected in this data set. Moreover, proliferation markers were expressed at similar levels in the mCherry-high/-negative cells, as observed in human cerebral organoids (Fig. S5C).

An expected enrichment of transcripts for *Tbr1* (5.97 log2FC), a layer 6 marker, and *Pcp4* (1.73 log2FC) and *Bcl11b* (2.12 log2FC); layer 5 markers, was observed in *Neurog2-mCherry*-high cells (Fig. 6B). In contrast, most upper-layer markers were expressed at low levels, with the exception of *Pou3f2* (Fig. S5D). The enrichment of early born neuronal markers is consistent with E12.5 as the stage of analysis, when early-born neurons are generated.

Transcripts for pan-neuronal markers, including *Tubb3* (3.65 log2FC)*, Dll1* (2.87 log2FC)*, Stmn2* (4.65 log2FC), and *Gap43* (2.41 log2FC) were all also enriched in mCherry-high cortical cells (Fig. S5E). Notably, at this early stage of development, glial genes were not expressed at high enough levels to compare. Thus, although mCherry is preferentially enriched in IPC subpopulations in human and murine cortices, ECM gene expression is only elevated in mCherry-high cells from human cerebral organoids.

### Murine *Neurog2* induces *Ppp1r17* and *Col3a1* gene expression in P19 cells

*Neurog2* is expressed in the vast majority of basal NPCs during early murine corticogenesis (Moffat et al., 2023), yet ECM gene expression is not detected in the germinal compartments. However, NEUROG2 transcriptional activity is dosage sensitive (Kovach et al., 2013), with phosphorylation of SP/TP sites by proline-directed serine-threonine kinases (e.g., GSK3, CDKs) inhibiting the ability of NEUROG2 to transactivate its target genes (Ali et al., 2011; Hardwick et al., 2015; Li et al., 2012). Thus, we asked whether *Neurog2* was sufficient to induce ECM gene expression when overexpressed at supraphysiological levels, using a native *Neurog2* gene or a mutated version (*Neurog2*^SA9TA1^) containing nine serine-to-alanine and a threonine-to-alanine mutations. We specifically asked whether *Neurog2* and/or *Neurog2*^SA9TA1^ were able to induce the expression of ECM and OPC genes, as well as *Ppp1r17,* a critical regulator of the timing of human cortical neurogenesis. P19 cells were transfected with pCIG2-GFP, pCIG2-*Neurog2*-GFP or pCIG2-*Neurog2*^SA9TA1^-GFP expression vectors. After 2 days *in vitro*, the ability of *Neurog2* and *Neurog2*^SA9TA1^ to induce the expression of known target genes was first assessed by qPCR. *Neurog2*^SA9TA1^ was significantly better at turning on the expression of *Dll3, Rnd2* and *Neurod1* than native *Neurog2,* as anticipated (Fig. 6E). *Neurog2*^SA9TA1^ also induced the ectopic expression of *Ppp1r17*, a human IPC gene (Fig. 6F) and *Col3a1*, an ECM gene (Fig. 6G), at higher levels than *Neurog2.* However, the overexpression of *Neurog2* or *Neurog2*^SA9TA1^ could not induce OPC gene expression in P19 cells (Fig. 6H). Thus, even though the *Neurog2*^+^ murine cortical lineage does not normally express ECM genes or *Ppp1r17*, *Neurog2* is sufficient to induce the expression of at least a subset of these genes in murine P19 cells, especially when inhibitory phosphoacceptor sites are mutated.

## DISCUSSION

In this study, we engineered *NEUROG2-mCherry* KI reporter hESCs to facilitate cerebral organoid modeling of *NEUROG2* function during human cortical development. We found that in derivative organoids, *NEUROG2* expression is enriched in IPCs, basal NPCs that occupy an expanded oSVZ during human neocortical development. Using short-term lineage tracing, we further found that *NEUROG2* regulates a human-specific neurodevelopmental gene regulatory program. Key human-specific genes regulated by *NEUROG2* include *PPP1R17*, a phosphatase-encoding gene that slows down cell cycle progression and overall rates of neurogenesis (Girskis et al., 2021), and ECM remodeling genes, which drive basal progenitor cell expansion (Amin and Borrell, 2020; Arai et al., 2011; Fietz et al., 2012; Florio et al., 2015; Martinez-Martinez et al., 2016; Pollen et al., 2015). These findings provide important new insights into the molecular mechanisms that distinguish the gene regulatory pathways that control murine versus human neurogenesis, allowing for the formation of lissencephalic versus gyrencephalic cortices, respectively.

The ECM regulates tissue stiffness, creating biophysical signals that operate together with biochemical (*e.g.,* morphogens) cues in the microenvironment (*i.e.,* niche) to control NPC proliferation and differentiation (Chan et al., 2017; Chaudhari et al., 2014; Gattazzo et al., 2014; Guilak et al., 2009; Mohyeldin et al., 2010). These biophysical and biochemical signals differ across species and are continually remodeled as the brain develops and matures into adulthood (Nelson, 2009; Rozario and DeSimone, 2010). In rodent cortices, ECM expression levels are high in the VZ, where proliferative, aRG reside, but not in the SVZ, where IPCs have a limited proliferative potential (Arai et al., 2011; Florio et al., 2015; Pollen et al., 2015). In contrast, in gyrencephalic cortices, bRG and IPCs express high levels of ECM genes to support integrin signaling and create a pro-proliferative, oSVZ niche (Amin and Borrell, 2020; Arai et al., 2011; Fietz et al., 2012; Martinez-Martinez et al., 2016). As a result, bRG and IPCs, which have lost constraining attachments to the ventricular surface, proliferate extensively to support cortical folding. By demonstrating that *NEUROG2* is expressed in IPCs and regulates ECM gene expression, we have found a potential explanation for the expanded ECM production by basal NPCs during human cortical development.

The link between *NEUROG2-mCherry* expression and the oligodendrocyte lineage was also unexpected but may relate to the existence of Neurog2-Ascl1 co-expressing cortical NPCs in the rodent cortex. These cortical NPCs have a mixed potential to give rise to neurons or oligodendrocytes, as we previously demonstrated (Han et al., 2021). Indeed, OPC marker expression was observed in *Neurog2-mCherry* cortical cells from rodents at later stages of development (postnatal day – 2) (Han et al., 2021), which is the period of oligodendrocyte differentiation in rodents (Kessaris et al., 2006). In gyrencephalic cortices, neurogenic and oligodendrogenic NPCs co-exist at the same time during the expansion phase of the oSVZ (Martinez-Cerdeno et al., 2012; Rash et al., 2019; Zecevic et al., 2005). In this study, we showed that Neurog2 regulates ECM production, and ECM molecules promote OPC proliferation (Hu et al., 2009), suggesting that NEUROG2 may support the expansion of OPCs in an autocrine fashion. SOX9 is similarly expressed in basal NPCs in the human and ferret cortex, and its overexpression in murine basal NPCs induces ECM gene expression, downregulates Eomes, and induces the expression of gliogenic genes such as Olig2 (Guven et al., 2020). The relationship between SOX9 and NEUROG2 during ECM deposition in human cortical development will be an area of future study.

A marked number of new genes have evolved to drive human brain development and functioning, originating either via gene duplications, structural/functional alterations of ancestral genes or their gene regulatory regions, or from *de novo* origins (Hodge et al., 2019). Our study suggests that human *NEUROG2* has acquired new target genes that contribute to the diversification of the developmental trajectories of cortical neural stem and progenitor cells. Altered target selection may arise due to modifications to the regulatory regions of target genes, new protein-protein interactions that change NEUROG2 activity, or other post-translational alterations. However, alterations to the core *NEUROG2* coding sequence seem unlikely to be the central driver of target switching. Indeed, human NEUROG2 protein has 84.2% homology with murine NEUROG2, with 99% homology in the bHLH domain, in keeping with the high degree of conservation of bHLH activity across invertebrate and vertebrate species (Guillemot and Hassan, 2017). Such conservation means that bHLH genes can compensate for each other to some extent. For instance, even though *Neurog2* is normally expressed in the dorsal telencephalon (i.e., the neocortical anlage), *Neurog2* can rescue ventral telencephalic neurogenesis defects when knocked-into the *Ascl1* locus (Parras et al., 2002). Moreover, when expressed at physiological levels with this knock-in approach, *Neurog2* does not respecify ventral NPCs, which still acquire a GABAergic identity (Parras et al., 2002). However, when *Neurog2* is overexpressed via electroporation under the control of a strong CAGG promoter, the supraphysiological levels allow *Neurog2* to respecify ventral telencephalic NPCs to a dorsal, glutamatergic identity (Kovach et al., 2013; Mattar et al., 2008). Thus, the proneural genes are dosage sensitive, raising the possibility that levels of expression distinguish target gene selection in murine and human cortical NPCs.

Proneural transcription factors induce neural stem cells to differentiate and sustain neural lineage fidelity in two ways: they directly transactivate downstream genes and they act as pioneer transcription factors, meaning they bind target sites in ‘closed’ chromatin and unwind tightly packed nucleosomes so other neurogenic transcription factors can bind (Aydin et al., 2019; Chanda et al., 2014; Fernandez Garcia et al., 2019; Park et al., 2017; Păun et al., 2023; Raposo et al., 2015; Smith et al., 2016; Soufi et al., 2015; Wapinski et al., 2017; Wapinski et al., 2013). By altering chromatin accessibility, proneural transcription factors may be central architects of the widespread epigenomic remodeling that occurs as NPCs traverse neurodevelopment and adulthood (Yang et al., 2023). Since pioneer transcription factor activity can be facilitated by interactions with chromatin remodeling complexes (Minderjahn et al., 2020) (*e.g.,* ASCL1 protein interacts with chromatin remodelers) (Păun et al., 2023) or by interactions with transcription factor partners (Cernilogar et al., 2019), changes to binding partners may account for differences in how murine and human NEUROG2 operate. Endogenous *Ascl1* has bona fide pioneer functions *in vivo* (Păun et al., 2023), but *Neurog2* has only been studied via over-expression (Aydin et al., 2019). Since over-expressed TFs can gain non-physiologic pioneer activity (Hansen et al., 2022), it is important that future studies examine pioneering functions of endogenous proneural transcription factors. The *NEUROG2-mCherry* KI hESCs generated in this study will facilitate this future area of study.

We and others found that proneural transcription factors are inhibited by phosphorylation in embryonic mouse brain and spinal cord, in Xenopus and fruit fly primary neurogenesis, and in cancer cells (Ali et al., 2011; Ali et al., 2020; Azzarelli et al., 2015; Azzarelli et al., 2022; Ge et al., 2006; Hand et al., 2005; Hindley et al., 2012; Li et al., 2014; Li et al., 2012; Quan et al., 2016; Sun et al., 2001). We used this knowledge to mutate *Ascl1* so that it cannot be phosphorylated and remains active, and showed that this designer proneural transcription factor is more efficient at lineage conversion of an astrocyte to an induced neuron than native *Ascl1* (Ghazale et al., 2022). In this study, we used a similar strategy to mutate SP and TP sites in *Neurog2* to generate *Neurog2*^SA9TA1^, and we demonstrated that this mutated version has an enhanced capacity to transactivate known *Neurog2* target genes, *Ppp1r17* and *Col3a1.* Thus, target gene selection may be differentially modulated in human versus rodent cortical NPCs depending on phosphorylation status of NEUROG2. Intriguingly, SOX9 is similarly expressed in basal NPCs in the human and ferret cortex, and its overexpression in murine basal NPCs induces ECM gene expression cell autonomously and non-autonomously (Guven et al., 2020). Thus, NEUROG2-expressing basal NPCs may be similarly influenced by signals emitted by other cells in the environment that alters, for example, phosphorylation status.

Taken together, these findings reveal that human *NEUROG2* and murine *Neurog2* have distinct transcriptional targets and altered functions during neocortical development. Future studies will be aimed at more comprehensively elucidating human-specific alterations to *NEUROG2* regulation and selection of downstream transcriptional targets during human neocortical development.

## MATERIALS AND METHODS

### Animal sources and maintenance

All animal procedures were performed with the approval of the SRI Animal Care Committee (AUP 21-769) and followed the Canadian Council on Animal Care guidelines. Animals were housed in the Sunnybrook Research Institute (SRI) Comparative Research Unit (CRU, SRI, Toronto, ON, Canada) with *ad libitum* access to food and water. *Neurog2^Flag-mCherry^*^KI/+^ mice were maintained on a CD1 background and genotyped as described (Han et al., 2021). The day of the vaginal plug was considered E0.5 for embryonic staging.

### hESC maintenance

Human embryonic stem cells (hESCs) (H1/WA01) were purchased from WiCell Research Institute, Wisconsin, USA. hESC usage for this project was approved by the Canadian Stem Cell Oversight Committee (SCOC application to CS and to CS and JN) as well as by SRI’s Research Ethics Board (REB Project Identification Number: 5003). Briefly, hESCs were cultured under feeder-free conditions in mTeSR Plus media (Stem Cell Technologies, 100-0276) on plates coated with Matrigel (Corning, 354277) and maintained in 5% CO_2_ incubators at 37°C. Versene (Thermo Fisher Scientific, 15040-066) was used to dissociate hESCs by manual pipetting every 4-5 days for maintenance. hESC cultures were monitored daily for differentiated cells, which were removed by manual scraping. Prior to generating cerebral organoids, quality control tests were routinely performed on hESC cultures, using a human stem cell pluripotency detection qPCR kit (Sciencecell, 0853) and hPSC genetic analysis kit (Stem Cell Technologies, 07550), according to the manufacturer’s protocols.

### CRISPR-Cas9 Gene-Editing

Clustered regularly interspaced short palindromic repeats (CRISPR) genome editing was used to insert an mCherry reporter gene into the 3’untranslated region (UTR) of the *NEUROG2* locus. The vector pSpCas9(BB)-2A-GFP (PX458) (Addgene, Plasmid 48138), was purchased to target NEUROG2. To promote homology-directed repair of the Cas9-cleaved *NEUROG2* target locus, we co-electroporated: (1) a CRISPR plasmid containing SpCas9-2A-eGFP and a single guide RNA (sgRNA) to *NEUROG2* that was a fusion of a CRISPR RNA (crRNA) and a trans-activating crRNA (tracrRNA); and (2) a repair template containing homology arms flanking an mCherry reporter cassette. 1.5 x 10^6^ cells were transfected with 1 µg DNA (500 ng Cas9 plasmid and 500 ng linearized donor plasmid) by nucleofection (pulse code CA137) using P3 Primary Cell 4D-Nucleofector X kit (Lonza, V4XP-3024) in a 4D-Nucleofector (Lonza, AAF-1003B) and were plated in a 6-well plate containing 10 µM Rock inhibitor Y-27632 (Stem Cell Technologies, 72302). Twenty-four hours post transfection, cells were harvested for quantification of transfection efficiency by flow cytometry for EGFP expression, indicative of Cas9 transfection. To isolate individual clones, positive GFP cells were sorted and replated at clonal density in multiple 10-cm plates. Each clone was expanded for 7-10 days and frozen while some cells were lysed for gDNA extraction for HDR screening with digital droplet PCR (ddPCR).

Individual clones were then picked and further expanded, followed by genomic DNA extraction to screen for clones that underwent homology-directed repair (HDR) using ddPCR. For ddPCR, we employed a forward primer-probe upstream to the starting point of the homology arm region and a reverse primer-probe that only bound to a site inside the exogenous mCherry sequence. The selected two correctly targeted clones were used for downstream experiments.

### Digital Droplet PCR

QX200 droplet digital PCR (ddPCR) system (Bio-Rad) was used for all ddPCR reactions. Detailed information for all primers is in Table S10. For HDR screening, the absolute number of *NEUROG2-mCherry* KI gene copies per cell was quantified and normalized to *RPP30* (Bio-Rad, 10031243). 20 ng of genomic DNA was used in 20 µl PCR reaction containing 900 nM of the forward and reverse *NEUROG2-mCherry* KI and *RPP30* primers, 250 nM of *NEUROG2-mCherry* KI and *RPP30* probes, and 10 µl of 2X ddPCR supermix for probes (Bio-Rad). Assay mixtures were loaded into a droplet generator cartridge (Bio-Rad), followed by the addition of 70μL of droplet generation oil for probes (Bio-Rad) into each of the eight oil wells. The cartridge was then placed inside the QX200 droplet generator (Bio-Rad). Generated droplets were transferred to a 96-well PCR plate (Eppendorf), which was heat-sealed with foil and placed in C1000 Touch Thermal Cycler (Bio-Rad). Thermal cycling conditions were as follows: 95°C for 10 min, 44 cycles of 94°C for 30 s, 53°C for 1 min, and 98°C for 10 min. FAM fluorescent signal, which labeled the *NEUROG2-mCherry* KI DNA sequence, and HEX fluorescent signal which labeled the *RPP30* DNA sequence, were counted by a QX200 digital droplet reader and analyzed by QuantaSoft analysis software ver.1.7.4.0917 (Bio-Rad). Identified positive clones were expanded and underwent further quality checks.

To quantify the absolute number of *mCherry* and *COL1A2* transcripts, RNA from mCherry-high and mCherry-low cell populations were collected from cerebral organoids. 10ng of total RNA was reverse-transcribed in a 10μL reaction using the SuperScript VILO cDNA Synthesis Kit (Invitrogen). The resulting cDNA was diluted to either 1:5 (*mCherry*) or 1:1500 (*COL1A2*) before amplification. The ddPCR reaction was performed in a 20μL volume containing 10μL of 2X QX200 ddPCR EvaGreen Supermix (Bio-Rad), 5μL of diluted cDNA, and 1μL each of 4μM forward and reverse primers and 3 μL of nuclease-free water. Droplet generation was completed as above but with the addition of 70μL of droplet generation oil for EvaGreen (Bio-Rad). Thermal cycling conditions were as follows: 95°C for 5 minutes, then 44 cycles of 96°C for 30 seconds and 56°C (*mCherry*) or 60°C (*COL1A2*) for 1 minute, and then 4°C for 5 minutes, 90°C for 5 minutes and 4°C for indefinite hold for dye stabilization. EvaGreen fluorescent signal in each droplet were counted and analyzed as described. The copy number of *mCherry* and *COL1A2* transcripts were normalized to the copies per ng of total RNA. All ddPCR analyses were performed at the SRI Genomics Core Facility.

### Cerebral organoid generation with dual SMAD inhibitors

We adapted our cerebral organoid differentiation protocol according to a previously described protocol (Qian et al., 2018; Qian et al., 2016). For embryoid body (EB) formation on day 0, hESC colonies were dissociated with Gentle cell dissociation reagent (GCDR, Stem Cell Technologies, 07174) for 7 mins at 37 °C and 12,000 cells in 100 µl STEMdiff kit EB formation media (Stem Cell Technologies, 08570) supplemented with 50 µM Rock inhibitor Y-27632 (Stem Cell Technologies, 72302) were plated in 96 well V-bottom plate (low-binding) (Greiner bio-one, 651970). On day 1 and 3, media was added with 2 µM Dorsomorphine, an inhibitor of BMP type I receptors (ALK2, ALK3, ALK6), (Stem Cell Technologies, 72102) and 2 µM A83-01, an inhibitor of TGFβ type I receptors (ALK4, ALK5, ALK7) (Stem Cell Technologies, 72022) for 5 days *in vitro* to induce embryoid body (EB) formation. On day 5, single EBs that reached ∼ 400-600 um diameter were selected for neural induction and were transferred to individual wells of 24-well ultra-low attachment plate (Corning, 3473) and media was replaced with STEMdiff kit Induction media consisting 1 µM SB431542, an ALK4, ALK5, ALK7 inhibitor (Stem Cell Technologies, 72234) and 1 µM CHIR99021, a GSK3β inhibitor that activates Wnt signaling and limits apoptosis (Delepine et al., 2021; Qian et al., 2016) (Stem Cell Technologies, cat. no. 72054) and cultured 4 more days. On day 9, EBs of ∼500-800 um diameter that had translucent edges, a sign of neuroepithelial induction were placed onto a single dimple on embedding sheet (Stem Cell Technologies, cat. no. 08579) and 15 µl of Matrigel, an undefined ECM preparation that contains collagens, laminins, other ECM molecules and growth factors (Corning, cat. 354277) was added to each cerebral organoid to encapsulate it. The Matrigel droplets were incubated at 37 °C for 30 mins before they were washed into 6-well ultra-low attachment plate (Stem Cell Technologies, 38071) containing STEMdiff kit Expansion media with 1 µM SB431542 and 1 µM CHIR99021. On day 13, individual EBs with clear neuroepithelial cell buds were transferred to each well of a 12-well miniature spinning bioreactor (Qian et al., 2018; Qian et al., 2016) containing STEMdiff kit Maturation media. From day 30, extracellular matrix (ECM) proteins were supplemented in Maturation media by dissolving Matrigel at 1% (v/v) containing human recombinant brain-derived neurotrophic factor (BDNF; PeproTech, AF-450-02). The aggregated cells were referred to as cerebral organoids from this stage onward and were allowed to further develop in maturation media until the experimental endpoints, as described.

### Cryosectioning and immunostaining

Cerebral organoids were rinsed with ice-cold phosphate-buffered saline (PBS, without Ca^2+^ and Mg^2+^) (Wisent, 311-010-CL), fixed in 4% paraformaldehyde in PBS (PFA, Electron Microscopy Sciences, 19208) overnight, and immersed in 20% sucrose (Sigma, 84097)/1X PBS overnight after three washes for 5 mins in PBS. Embryonic rodent brains were dissected from the skull before following the same procedures. Tissues were embedded in optimal cutting temperature compound (OCT), and 10-micron sections were collected with a Leica CM3050 cryostat (Leica Microsystems Canada Inc., Richmond Hill, ON, Canada). Samples were collected on Fisherbrand^TM^ Superfrost^TM^ Plus Microscope Slides (Thermo Fisher Scientific, 12-550-15). Cryosections of fixed cerebral organoids were washed in 0.1% Triton X-100 (Sigma, T8787) in PBS (PBST), then blocked for 1 h at room temperature in 10% horse serum (HS, Wisent, 065-150) in PBST. Primary antibodies were diluted in blocking solution as follows: SOX2 (1:500, Abcam #ab97959), PAX6 (1:500, Biolegend #901301), NEUROG2 (1:500, Invitrogen #PA5-78556), NESTIN (1:500, R&D Systems #MAB1259), DCX (1:500, Abcam #ab6586), COL4 (1:200, Abcam #ab6586), LAM (1:200, Sigma #L9393), mCherry (1:500, Sicgen #AB0040-200) and TUJ1 (1:500, BioLegend #802001). After one hour blocking at room temperature, slides were incubated with primary antibodies at 4℃ overnight. The next day, slides were washed 5 times for 5 mins in PBST, followed by incubation with 1:500 dilutions of species-specific secondary antibodies (Invitrogen Molecular Probes) for 1 h at room temperature. Slides were washed five times in PBST and counterstained with 4’,6-diamidino-2-phenylindole (DAPI, Invitrogen, D1306) and mounted in Aqua-polymount (Polysciences Ince., 18606-20). All images were taken using a Leica DMi8 Inverted Microscope (Leica Microsystems CMS, 11889113).

### Tissue clearing, immunolabeling, and lightsheet fluorescence microscopy

hESC-derived cerebral organoids at 42 and 93 days old were fixed in 4% PFA overnight at 4°C. To preserve the tissue protein architecture, samples were cleared using the SHIELD (Stabilization to Harsh conditions via Intramolecular Epoxide Linkages to prevent Degradation) method (Park et al., 2018). Specifically, organoids were incubated in SHIELD OFF solution at 4℃ with shaking for 24hrs. Subsequently, they were incubated in a mixture of SHIELD ON-Buffer and the SHIELD-Epoxy solution (7:1 ratio) at 37℃ with shaking for 6 hrs. Lastly, the samples were incubated in SHIELD ON-Buffer at 37°C with shaking overnight. To carry out tissue delipidation, the samples were passively run down in the Delipidation buffer (Lifecanvas Technologies) for 3 days at room temperature (RT). Samples were washed in 0.1% PBST 3 times over a span of 3 hours following each incubation and PFA fixation. Organoids were incubated in primary antibodies diluted in 0.1% PBST (see above) at RT for 48hrs. Samples were then incubated in conjugated secondary antibodies diluted in 0.1% PBST (Alexa Fluor 488 goat anti-mouse IgG2a, Alexa Fluor 568 donkey anti-rabbit, Alexa Fluor 488 donkey anti-rabbit IgG(H+L); 1:250) at RT for 48hrs. For index matching and to make samples optically transparent they were incubated in the EasyIndex medium (Lifecanvas Technologies, RI= 1.52) at RT overnight. Cerebral organoid images were acquired using the UltraMicroscope Blaze lightsheet fluorescence microscope (Miltenyi Biotech) with a 4X objective. The samples were mounted on a small sample stage using photoactivated adhesive (Bondic CNA) and placed into a custom organic imaging medium in the microscope’s chamber (Cargille Immersion Liquid, RI= 1.52). Two channels were acquired with a 488 nm wavelength and 85 mW power, and 639nm with 70 mW, for mCherry and SOX2/TUJ1/LAM or COL4, respectively. A 1.67x magnification post-objective lens was employed generating an in-plane resolution of 1.95 μm and a step size of 3.55 μm (scanning protocol parameters: laser sheet thickness = 7.1 μm, NA=0.050, and laser width = 30% single sided multi-angle excitation).

### Bulk RNA-seq

Cerebral organoids were dissociated into a single-cell suspension using Worthington Papain System kit (Worthington, LK003150) according to the manufacturer’s instruction. We collected single-cell suspensions from organoids at 45-47 days old and multiple organoids (7-8) were pooled together per sample to obtain a sufficient number of cells. Briefly, prewarmed papain solution with DNase (2.5 ml) was added to the organoids in a 60mm dish. Cerebral organoids were minced with a sterile razor blade into smaller pieces and incubated for 30-45 mins at 37°C on an orbital shaker (70 rpm). The tissue suspension was triturated 8-10 times with P1000 pipette tip to assist the release of single cells. Cell suspension was transferred to a 15 ml centrifuge tube and added with ovomucoid protease inhibitor solution to stop papain activity. The cells were centrifuged at 300g for 7 mins and filtered through 40 µm strainer to remove remaining cell aggregates. Single cells were resuspended in PBS containing FACS buffer (1 mM EDTA, 0.1% BSA and Ca^2+^/Mg^2+^ free PBS) with DAPI before sending for flow cytometry and each sample was sorted into mCherry-positive and mCherry-negative groups. Total RNA was extracted from FACS-isolated cells using MagMAX-96 total RNA isolation kit (ThermoFisher, AM1830). The extracted total RNA was quantified by Qubit 3 Fluorometer with Qubit RNA HS Assay kit. The integrity of total RNA (RIN value) was measured by Agilent 2100 Bioanalyzer with RNA 6000 Pico kit.

### Targeted transcriptome analysis

The targeted transcriptome sequencing was performed on the Ion S5XL Next Generation Sequencing system with the Ion AmpliSeq Transcriptome Human Gene Expression Assay (ThermoFisher Scientifics Inc). This assay covers 20,802 human RefSeq genes (>95% of UCSC refGene) with a single amplicon designed per gene target. The gDNA in the RNA sample was digested by ezDNase and the cDNA was synthesized from 10ng of total RNA using SuperScript IV VILO Master Mix with ezDNase Enzyme kit (Thermofisher Scientifics). The cDNA libraries were constructed by Ion Ampliseq Library Kit Plus. The targeted areas were amplified by polymerase chain reaction for 12 cycles. The resulting amplicons were treated with FuPa reagent to partially digest primers. Amplicons were ligated to Ion P1 and IonCode barcode adapters and purified using Agencourt AMPure XP reagent (Beckman Coulter). Barcoded libraries were quantified using the Ion Library TaqMan Quantitation Kit (ThermoFisher Scientific Inc) and diluted to a final concentration of 80pM. The sequencing template preparation was done using Ion Chef with Ion 540 Chef Kits. Sequencing was performed for 500 flows on an Ion S5XL Sequencer with Ion 540 chip.

### Next generation sequencing data analysis

The Ion Torrent platform-specific pipeline software, Torrent Suite version 5.18.1 (Thermo Fisher Scientific Inc) was used to separate barcoded reads and to filter and remove polyclonal and low-quality reads. Ion Torrent platform-specific plugin, ampliseqRNA (v5.18.0.0) was used for the alignment of the raw sequencing reads and quantitation of normalized gene expression level (reads-per-million:RPM). DESeq2 was used to do differential expression. Principal component analysis, generation of scatter plot and volcano plot, hierarchical clustering and pathway analysis were performed by Transcriptome Analysis Console (TAC) 4.0 software using the CHP files.

### Single cell RNA-seq data analysis in human fetal cortices and cerebral organoids

A Seurat v.3.2.3 R package was used to analyze the scRNA seq data in human cerebral organoids available from GSE137877 (Sivitilli et al., 2020). The scRNA seq data of human fetal cortices was available from GSE104276 (Zhong et al., 2018). Cells with more than 4000 genes and less than 500 genes were filtered out as doublets or low-quality cells, respectively. SCTransform function was used for data transformation followed by clustering with RunPCA, FindNeighbours and FindClusters, using the first 30 principal components. RunUMAP function was used to generate 2D projections of cell clustering, and cell type annotation was done using the described genes (Sivitilli et al., 2020). Proneural negative, *NEUROG2* or *NEUROG1* single and double positive cells were identified with an expression threshold greater than 0. Monocle3 R package was used for a pseudotime analysis and differentially expressed genes (DEGs) were reported using differential Gene Test function with an adjusted p value less than 0.001.

### P19 cell transfection

P19 cells were transfected using Lipofectamine 3000 with pCIG2, pCIG2-Neurog2 and pCIG2-Neurog2SA9TA1 DNA. The cell growth media was changed to fresh media 24 hrs post transfection and the cells were harvested 48 hrs post transfection.

### RNA isolation, cDNA preparation and qPCR

Total RNA extraction was performed from P19 cells using Qiagen RNA isolation mini kit (#74104), followed by CDNA preparation using RT2 First strand reverse transcription kit (#330401). qPCR using specific primers targeting known and predicted targets of *Neurog2* was performed using RT2 SYBR green qPCR Kit (#330513). The Qiagen primers used were *Dll3* (#PPM25734G-200), *Neurod1* (#PPM05527D-200), *Col1a1* (#PPM03845F-200), *Col3a1* (#PPM04784B-200), *Sox10* (#PPM04723A-200) and *Pdgfra* (#PPM03640D-200). qPCR analysis was performed using 2^-ΔΔCT method by normalizing to the CT value of pCIG2 control. This experiment was repeated 3 independent times (N=3 biological replicates) with 3 technical replicates per sample in each experiment. The trend was the same in all experiments. A single representative result is shown.

### Imaging, quantification, and statistics

Imaging was performed using epifluorescence microscope followed by analysis and quantification using Photoshop. Statistical analysis was performed using GraphPad Prism software.

## DATA AVAILABILITY

We mined scRNA seq data in human cerebral organoids available from GSE137877 (Sivitilli et al., 2020) and scRNA seq data of human fetal cortices was available from GSE104276 (Zhong et al., 2018). RNA-seq data from this study is presented in Supplemental Table 2.

## ACKNOWLEDGEMENTS

The authors acknowledge the support of the Sunnybrook Research Institute (SRI) Genomics Core Facility, the SRI Histology Core Facility (Petia Stefanova), and the Sunnybrook Centre for Cytometry and Scanning Microscopy (Kevin Conway). We also acknowledge The Imaging Facility at The Hospital for Sick Children for assistance with lightsheet microscopy.

## DATA AVAILABILITY

All relevant data can be found within the article and its supplementary information.

